# Spinning sugars in antigen biosynthesis: a direct study of the *Coxiella burnetii* and *Streptomyces griseus* TDP-sugar epimerases

**DOI:** 10.1101/2021.10.25.465559

**Authors:** Alice R. Cross, Sumita Roy, Mirella Vivoli Vega, Martin Rejzek, Sergey A. Nepogodiev, Matthew Cliff, Debbie Salmon, Michail N. Isupov, Robert A. Field, Joann L. Prior, Nicholas J. Harmer, on behalf of the GoVV consortium

## Abstract

The sugars streptose and dihydrohydroxystreptose (DHHS) are unique to the bacteria *Streptomyces griseus* and *Coxiella burnetii* respectively. Streptose forms the central moiety of the antibiotic streptomycin, whilst DHHS is found in the O-antigen of the zoonotic pathogen *C. burnetii*. Biosynthesis of these sugars has been proposed to follow a similar path to that of TDP-rhamnose, catalysed by the enzymes RmlA/RmlB/RmlC/RmlD. Streptose and DHHS biosynthesis unusually require a ring contraction step that might be performed by the orthologues of RmlC or RmlD. Genome sequencing of *S. griseus* and *C. burnetii* proposed the StrM and CBU1838 proteins respectively as RmlC orthologues. Here, we demonstrate through both coupled and direct observation studies that both enzymes can perform the RmlC 3’’,5’’ double epimerisation activity; and that this activity supports TDP-rhamnose biosynthesis in vivo. We demonstrate that proton exchange is faster at the 3’’ position than the 5’’ position, in contrast to a previously studied orthologue. We solved the crystal structures of CBU1838 and StrM in complex with TDP and show that they form an active site highly similar to previously characterised enzymes. These results further support the hypothesis that streptose and DHHS are biosynthesised using the TDP pathway and are consistent with the ring contraction step being performed on a double epimerised substrate, most likely by the RmlD paralogue. This work will support the determination of the full pathways for streptose and DHHS biosynthesis.

## Introduction

The gammaproteobacterium *Coxiella burnetii* evolved recently from tick endosymbionts^1^ to become an obligate intracellular pathogen of mammals ^2^. This bacterium infects a wide range of mammals causing reproductive failures ^3,4^. It is a significant economic pathogen of sheep, goats and cattle ^5^, causing spontaneous abortion of pregnancies ^6^. Humans can be infected by *C. burnetii* either through aerosols or consumption of meat and dairy products (with 1-10 bacteria sufficient to cause disease) ^7,8^. *C. burnetii* in unpasteurised ruminant milk poses a particular risk ^9-11^. Humans generally present with a self-limiting febrile illness ^12^. A small percentage of cases lead to more serious complications such as miscarriage, hepatitis, or endocarditis ^2,13,14^. *C. burnetii* infection can be effectively treated with doxycycline ^2^. However, diagnosis is often challenging as *C. burnetii* grows only in highly defined media, requires environment-controlled microaerobic incubators, ^15^ and symptoms of infection are generally non-specific.

From a “One Health” perspective, a vaccine against *C. burnetii* would be the ideal approach to reduce veterinary and human infection ^6^. Inactivated whole cell vaccines have proved effective in animals and humans ^16,17^. These are not licensed for human use in the USA, EU or UK as they cause severe reactions in seropositive individuals ^18^. Current efforts to develop novel vaccines are focused on development of subunit ^19^ or epitope based vaccines ^20,21^. A very strong subunit vaccine candidate is the *C. burnetii* polysaccharide O-antigen. A complete O-antigen is required for immune evasion and efficient infection of mammalian cells ^22,23^. Serial passaging of *C. burnetii* selects for mutants that cease producing some or all their O-antigen ^24,25^. The most common mutants either lack a polymer of the unusual saccharide virenose (“intermediate”), or the entire O-antigen ^26^ (Figure 1A). Strains lacking the O-antigen are avirulent in the guinea-pig model of infection ^27^. The O-antigen is the dominant epitope of wild-type *C. burnetii* ^28^, suggesting that any vaccine against *C. burnetii* will require the O-antigen ^28^. This highlights the importance of a deepened understanding of the biosynthesis of this polysaccharide.

**Figure 1.**
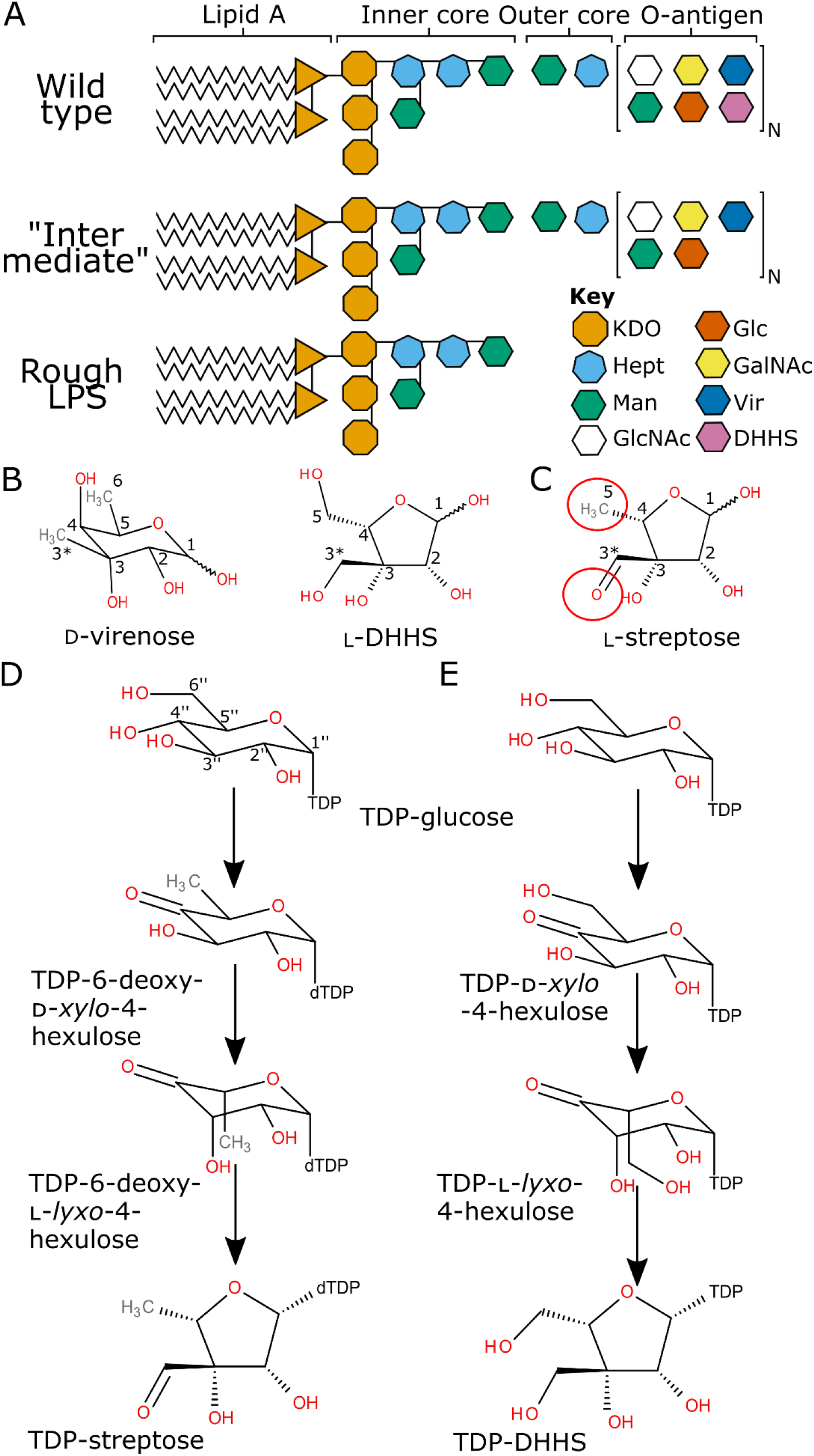
Saccharide structures in *Coxiella burnetii* and related sugars. **A**. Overview of the *C. burnetii* lipopolysaccharide and O-antigen. Wild-type *C. burnetii* makes a smooth LPS with the inner and outer cores and a full O-antigen. Mutants cause either an “intermediate” phenotype without virenose in the O-antigen, or a “rough” phenotype lacking the outer core and O-antigen. The linkage of the outer core and O-antigen is not known. Key abbreviations: KDO: 2-keto-3-deoxy-D-mannoctanoic acid; Man: mannose; GlcNAc: *N*-acetyl-glucosamine; Glc: glucose; GalNAc: *N-*acetyl-galactosamine; Vir: virenose; DHHS: dihydrohydroxystreptose. Based on ^26^. **B, C**. The unusual monosaccharides virenose and dihydrohydroxystreptose (DHHS) from the *C. burnetii* O-antigen (**B**) and the related sugar streptose (**C**). The differences between DHHS and streptose are highlighted with red circles. **D, E**. Proposed biosynthetic pathways for TDP-linked streptose (**D**) and DHHS (**E**). Chemical structures drawn using BIOVIA Draw v16.1.

The *C. burnetii* O-antigen characteristically contains two C-3 methylated/hydroxymethylated sugars, virenose and dihydrohydroxystreptose (DHHS; Figure 1B) ^29,30^. Virenose is also found in *Bacillus cereus* and *Streptomyces albaduncus* ^31,32^. Biosynthetic pathways have been proposed in these two organisms, and a similar pathway has been proposed in *C. burnetii* ^33^. No equivalent pathway has been proposed for DHHS biosynthesis. The most similar known sugar to DHHS is streptose (Figure 1B), the central saccharide unit of the aminoglycoside antibiotic streptomycin ^34^, produced by *Streptomyces griseus*. Streptose differs from DHHS in having no hydroxyl at C-6; and being oxidised at the C-3 hydroxymethyl group. Streptose is synthesised from TDP-glucose, in a manner analogous to the biosynthesis of rhamnose (Figure 1C) ^34-37^. A reasonable path to DHHS synthesis would involve a C-4 oxidase of TDP-glucose, a 5’’-mono or 3’’,5’’-double epimerase ^38^, and an enzyme similar to the streptose synthase ^37^ (Figure 1D,E). Although it remains possible that the epimerase might also perform the ring rearrangement, the only precedent for this reaction was performed by an enzyme from a different structural class ^39^.

Two operons (*cbu0825* to *cbu0856*, and *cbu1831* to *cbu1838*) have been suggested as coding for DHHS synthesising genes ^26^. The *cbu1831* to *cbu1838* contains two genes that show high sequence identity to genes in the rhamnose biosynthesis pathway. CBU1834 has over 60% sequence identity to characterised RmlA orthologues ^33^, and so is highly likely to be a glucose-1-phosphate thymidylyltransferase. TDP-glucose has also been proposed to be a precursor of virenose ^33^, and so is not necessarily characteristic of DHHS biosynthesis. CBU1838 shows 48% identity with characterised RmlC orthologues. With this sequence identity, this protein is highly likely to be an orthologue of RmlC and act as a TDP-D-*xylo*-4-hexulose 3,5-epimerase. This activity is not required for virenose biosynthesis. As *C. burnetii* does not produce rhamnose, it is unlikely that it would be required for rhamnose biosynthesis. This leaves DHHS biosynthesis as the only reasonable function of CBU1838.

RmlC is part of a family of epimerases that use a deprotonation/reprotonation mechanism^40^. RmlC uses a histidine base to deprotonate the substrate, and a tyrosine acid to reprotonate from the opposite side of the sugar ^41-44^. The same residues are likely involved in epimerising at both 3’’ and 5’’ positions: the sugar ring flips between the two reactions to present the second site to be epimerised to the catalytic base and acid^41^. These amino acids are highly conserved in the family (Figure 2). RmlC orthologues have been characterised biochemically and structurally from a range of species (Table 1) ^38,42,43,45,46^. Paralogues that selectively epimerise at either the 3’ or 5’ carbon have also been characterised^44,47,48^. These studies have highlighted likely residues for binding to the TDP carrier^43,49,50^. For streptose and DHHS biosynthesis, only epimerisation at the 5’’ position is strictly necessary: the C3-C4 bond is broken in the streptose biosynthesis step^35^, which will remove the stereocentre at C3. RmlC paralogues for streptose (StrM) and DHHS (CBU1838) biosynthesis have been proposed bioinformatically^33^ (e.g. by KEGG ^51^); however, such assignments can be inaccurate and so experimental confirmation is desirable ^52^.

**Table 1:** Summary of existing structures of RmlC paralogues. Key published structures of proteins with experimentally determined activities. ^a^Species abbreviations: Mtu: *Mycobacterium tuberculosis*; Pa: *Pseudomonas aeruginosa*; Ss: *Streptococcus suis*; Mth: *Methanobacterium thermoautotrophicum*; Sty: *Salmonella typhimurium*; Ba: *Bacillus anthracis*; Ao: *Amycolatopsis orientalis*; Sb: *Streptomyces bikiniensis*; Sn: *Streptomyces niveus*; Ye: *Yersinia enterocolitica*; At: *Aneurinibacillus thermoaerophilus*; Tt: *Thermoanaerobacterium thermosaccharolyticum*; Cj: *Campylobacter jejuni*.

| Reaction | Enzyme activity | EC number | Common name | PDB ID | Species<sup>a</sup> | Ref. |
| --- | --- | --- | --- | --- | --- | --- |
| 1 | TDP-6-deoxy-D-xylo-4-hexulose 3'',5''-epimerase | 5.1.3.13 | RmlC | 2IXC/1UPI | Mtu | 41,68 |
|  |  |  |  | 2IXH | Pa | 41 |
|  |  |  |  | 1NYW | Ss | 50 |
|  |  |  |  | 1EP0 | Mth | 45 |
|  |  |  |  | 1DZR | Sty | 38 |
|  |  |  |  | 3RYK | Ba | 46 |
| 2 | TDP-6-deoxy-D-xylo-4-hexulose 5''-epimerase | N/A | EvaD | 1OI6 | Ao | 47 |
| 3 | TDP-6-deoxy-D-xylo-4-hexulose 3''-epimerase | 5.1.3.27 | ChmJ | 4HN0 | Sb | 44 |
|  |  |  | NovW | 2C0Z | Sn | 43 |
| 4 | CDP-6-deoxy-D-xylo-4-hexulose epimerase | N/A | WbcA | 5BUV | Ye | 48 |
| 5 | TDP-6-deoxy-3,4,keto-hexulose isomerase | 5.3.2.4 | FdtA | 2PA7 | At | 69 |
|  |  |  | QdtA | 4ZU7 | Tt |  |
|  |  |  | WlaRB | 5TPU | Cj | 70 |
| 6 | GDP-D-glycero-4-keto-D-lyxo-heptose-3'',5''-epimerase | 5.1.3 | Cj1430 | 7M15 | Cj | 42 |

**Figure 2:**
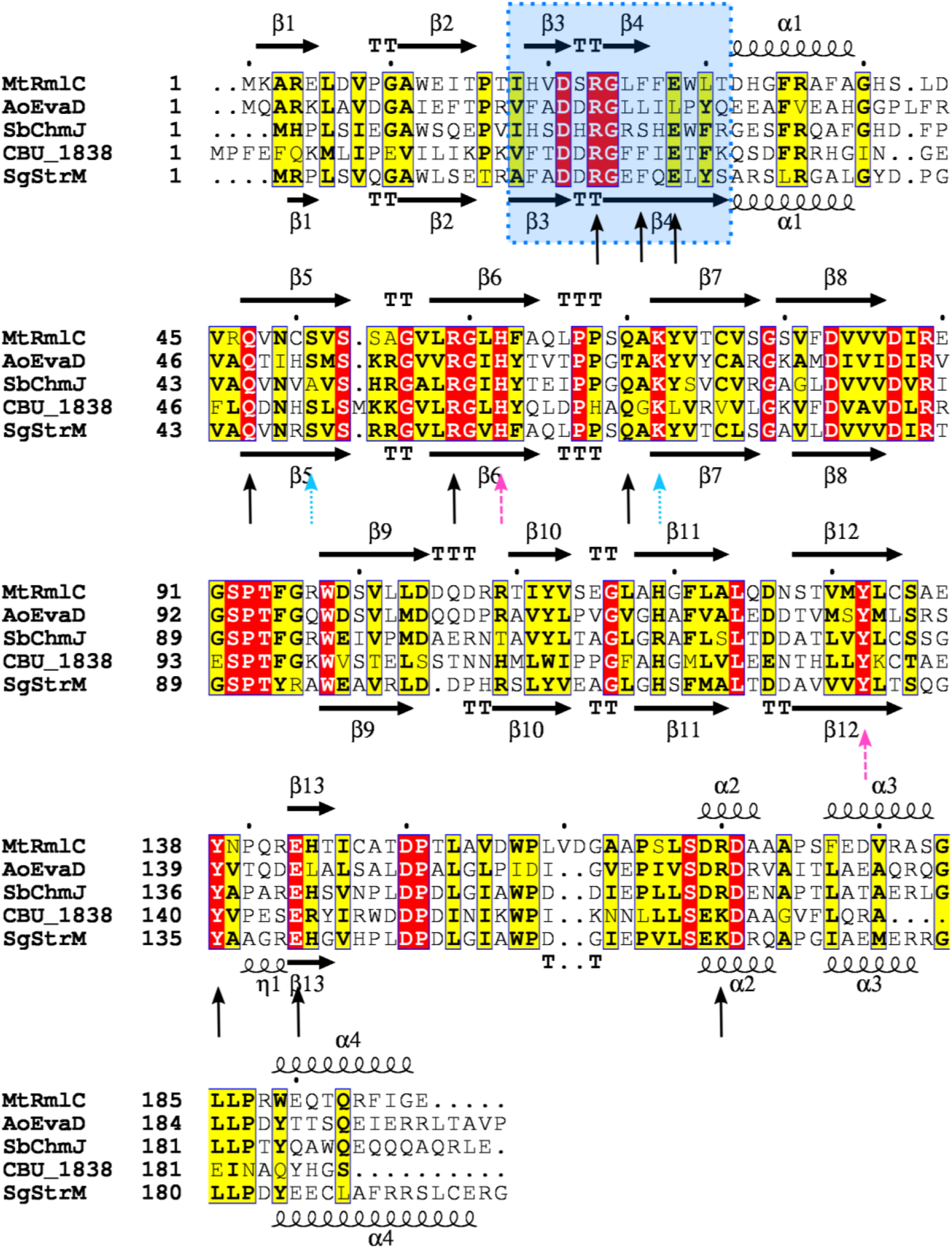
Structural alignment of RmlC paralogues. Alignment generated using Modeller v9.21 ^71^ from the PDB ID codes 2IXC ^41^, 1OI6 ^47^ and 4HMZ ^44^ (*Mt*RmlC, *Ao*EvaD and *Sb*ChmJ, respectively), and the PDB files from CBU1838 and SgStrM solved in this study. Chain A was used in each case. Secondary structure elements for *Mt*RmlC and *Sg*StrM are noted above and below the sequence alignments, respectively. The strand exchange between chains is highlighted with a blue box. The residues that interact with TDP and citrate, and which are likely the catalytic acid and base are indicated with black arrows, sky blue dotted arrows and pink dashed arrows respectively. Figure generated using ESPript v3.0222 ^72^; a red box/white character relates strict identity, a black bold character relates similarity in a group, and filled in yellow characters relates similarity across groups.

Here, we tested the hypothesis that StrM and CBU1838 are orthologues of RmlC. We expected that they would perform only the epimerisation step, and not perform the structural rearrangement activity that has yet to be assigned. As both enzymes contain all the catalytic residues proposed for 3’’,5’’-double epimerisation, we expected that they might perform both transformations. We demonstrate *in vivo* and *in vitro* that these enzymes are competent to perform the double epimerisation. Using NMR, we demonstrate that, surprisingly, proton-deuteron exchange at the 3’’ position is faster than at the 5’’ position for both enzymes. The structures of CBU_1838 and StrM show that they form an active site highly similar to previously characterised enzymes, in keeping with the biochemical data. These data support a 3’’, 5’’-double epimerised substrate for the streptose ring re-arrangement.

## Results

### CBU1838 complement RmlC in *E. coli*

We firstly tested whether CBU1838 can complement the *E. coli* RmlC orthologue RfbC. We cultured wild-type MG1655 *E. coli* and *rfbC* mutant bacteria, and *rfbC* mutants complemented with CBU1838. We extracted soluble small molecules from each culture. No TDP-linked sugars were detected in wild-type MG1655. In contrast, the *rfbC* mutant contained both TDP-glucose and TDP-6-deoxy-D-*xylo*-4-hexulose (Figure 3), as expected for a mutant at this essential step of the pathway. Neither of these compounds were detected when the *rfbC* mutant was complemented with CBU1838, restoring wild-type activity (Figure 3). These data gave good confidence that CBU1838 performs both 3’’ and 5’’ epimerisations of TDP-6-deoxy-D-*xylo*-4-hexulose, as expected.

**Figure 3:**
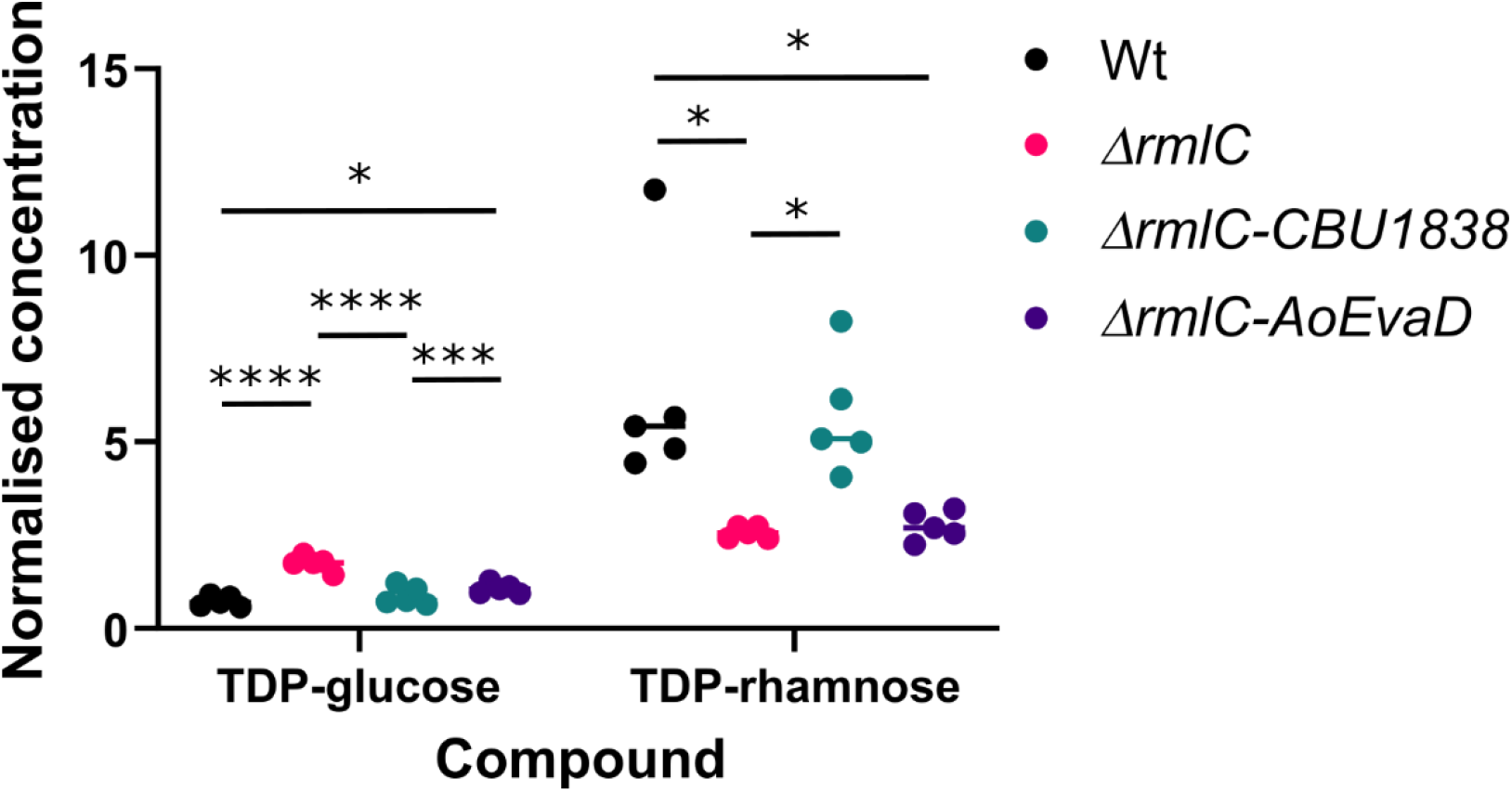
CBU1838 complements *rmlC* in vivo. MG1665 *E. coli*, the Keio collection *rmlC* mutant, and complements of this mutant with *CBU1838* and *AoEvaD* were grown to stationary phase in LB. Cells were harvested and metabolites extracted in 50% (v/v) acetonitrile. Metabolite concentrations were determined by HPLC-MS using a triple-quad instrument. Concentrations were determined by comparison to standards at defined concentrations and normalised to an internal ^13^C labelled TDP-glucose sample added to all samples. Five experimental replicates were taken for each data point. Bars represent the mean. All points for each compound were compared by ANOVA followed by Tukey’s multiple comparison test using Graphpad Prism v.9.2.0. ^*^: *p* < 0.05; ^***^: *p* < 0.0005; ^****^: *p* < 0.0001

### StrM and CBU1838 complement RmlC in vitro

We next tested whether StrM and CBU1838 can perform the function of *E. coli* RmlC in a coupled assay. We used the test enzyme coupled to RmlB and RmlD in a catalytic cascade to convert TDP-glucose to TDP-rhamnose. Both StrM and CBU1838 complemented RmlC in these assays effectively (Figure 4). Identical concentrations of StrM and RmlC gave a similar effect. A higher concentration of CBU1838 was required to achieve the same effect. This is consistent with the expected function of CBU1838, epimerising a sugar that retains the 6’’-OH group.

**Figure 4:**
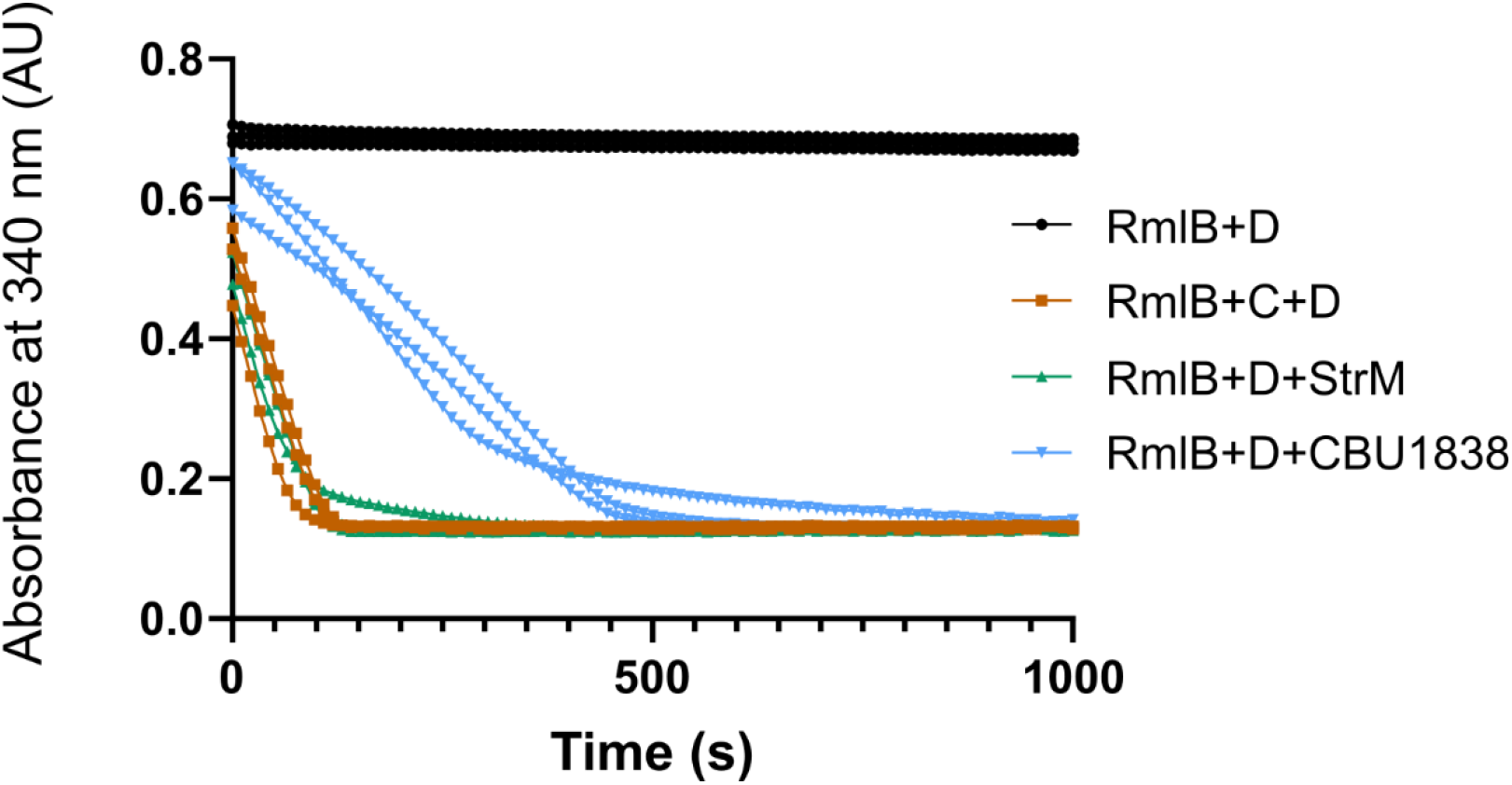
StrM and CBU1838 complement RmlC *in vitro*. The turnover of TDP-glucose to TDP-rhamnose by RmlB, RmlC and RmlD was monitored by following the reduction of NADPH at 340 nm. 200 M each NADPH and TDP-glucose were incubated with 0.4 M RmlB and RmlD, and as appropriate 0.4 M RmlC, 0.1 M StrM, or 1.6 M CBU1838. StrM and CBU1838 were able to complement RmlC in this assay, albeit at a reduced rate for CBU1838. Three experimental replicates were taken for each condition.

These data further support the concept that StrM and CBU1838 perform both 3’’ and 5’’ epimerisations of TDP-6-deoxy-D-*xylo*-4-hexulose.

### CBU1838 shows a reduced rate compared to RmlC and StrM, consistent with its modified substrate

We determined the kinetic constants for StrM and RmlC, in comparison to other characterised enzymes. We used purified TDP-6-deoxy-D-*xylo*-4-hexulose and coupled the test enzymes to RmlD. Data from StrM, RmlC, CBU1838 and EvaD all fitted very well to the Michaelis-Menten equation, with no evidence of cooperativity (Figure S1). StrM and RmlC showed a similar *k*_*cat*_, with StrM having a slightly lower *K*_*M*_ (Table 2). CBU1838 showed a lower *k*_*cat*_ and a clearly higher *K*_*M*_ (Table 2). This is consistent with its expected 6’’-hydroxy substrate. As expected, the 5’’-specific epimerase EvaD showed a substantially lower *k*_*cat*_. We did not observe any activity with the 3’’-specific epimerase ChmJ (Figure S2). Previous studies have demonstrated that ChmJ has no 5’’-epimerisation activity ^44^, and that the coupling enzyme RmlD only reduces double-epimerised products ^53,54^. The kinetic constants provide further evidence that StrM is likely an orthologue of RmlC. CBU1838 can performs this activity, but its likely native substrate is 6’’-hydroxylated.

**Table 2:**
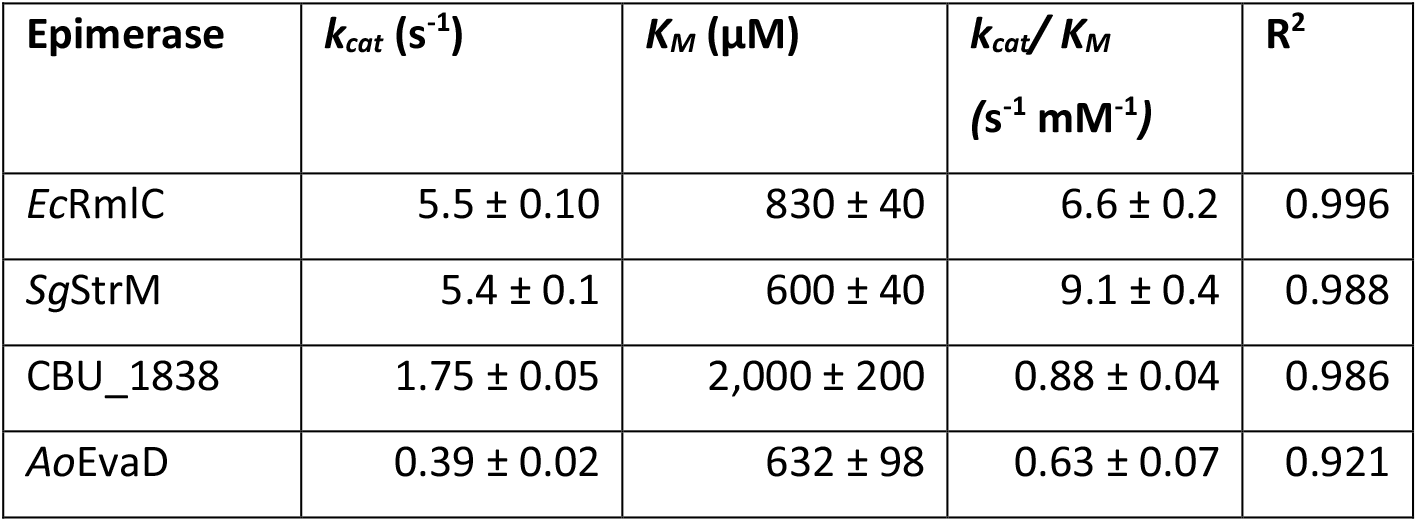
Kinetic constants for *Ec*RmlC, *Sg*StrM, *Cb*CBU_1838 and *Ao*EvaD. Kinetic data were determined using commercial TDP-6-deoxy-D-*xylo*-4-hexulose, with the reaction coupled to *Ec*RmlD to allow continuous measurement of the oxidation of NADPH. Data were fitted to the Michaelis-Menten equation using Graphpad v. 8.0 and the fit quality determined from the R^2^ value.

### RmlC, StrM and 1838 epimerise at both the 3’’ and 5’’ positions

To give further confidence in the activity of CBU1838 and StrM, in a novel assay, we monitored their activity directly using NMR and GC-MS without employing a coupling agent. We performed ^1^H-NMR experiments in deuterated buffer. As RmlC uses a deprotonation-protonation mechanism with different proton acceptors and donors, we expected to observe a loss of signal from protons bound to the 5’’ and 3’’ carbon atoms. We observed solvent exchange at both positions for RmlC, StrM, CBU1838 and EvaD, with relative rates in the same order as the rates observed in a coupled assay (Figure 5A, S3). Surprisingly, the rate of exchange at the 3’’ position was significantly faster than the rate at the 5’’ position for RmlC, StrM and CBU1838 (Table 3); for RmlD the rate of exchange at the two positions was similar. When the reaction was permitted to proceed to completion, RmlC, StrM and CBU1838 all showed almost complete exchange at both 3’’ and 5’’ positions (Figure 5B). In contrast, the 3’’ specific enzyme ChmJ showed almost complete exchange at the 3’’ position, with very little exchange observed at the 5’’ position. GC-MS analysis of enzyme reactions run to completion confirmed that exchange had occurred at both positions for CBU1838, StrM and RmlC (Figure S4). Consistent with the NMR data and previous work^44^, ChmJ showed strong exchange at the 3’’ position only. These data strongly support a double epimerase activity for both CBU1838 and StrM.

**Figure 5:**
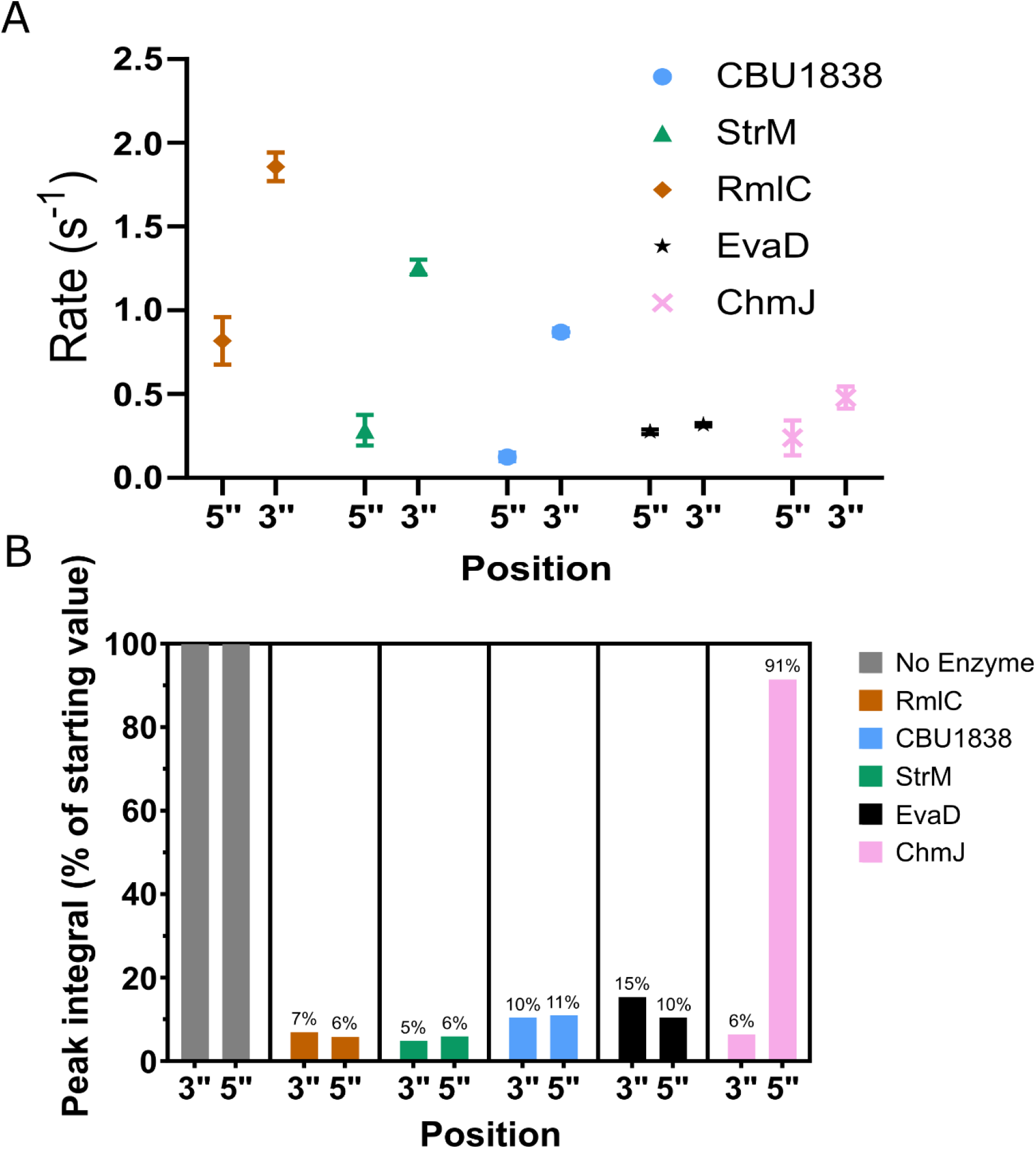
Rate of RmlC paralogues monitored by NMR. RmlC paralogues were incubated with purified TDP-6-deoxy-D-*xylo*-4-hexulose. **A**. The loss of proton signal at the 3’’ and 5’’ positions in the sugar was monitored by NMR. The rate was monitored every 5-10 minutes over 90 minutes to calculate an initial rate. **B**. Samples were incubated over 18 hours to allow the reaction to proceed to completion. RmlC, StrM and CBU1838 showed almost complete conversion to product in both 3’’ and 5’’ positions. ChmJ in contrast showed conversion only at the 3’’ position. Note that the substrate showed almost no loss of signal over the course of the experiment. Error bars represent the standard error in the mean.

**Table 3:** Rate of hydrogen-deuterium exchange catalysed by RmlC paralogues. RmlC paralogues were incubated with commercial TDP-6-deoxy-D-*xylo*-4-hexulose. The loss of proton signal at the 3’’ and 5’’ positions in the sugar was monitored by NMR. The rate was monitored for up to 90 minutes to calculate an initial rate.

| Epimerase | Rate of H-D exchange (s<sup>-1</sup>) |  |
| --- | --- | --- |
|  | 3'' | 5'' |
| **RmlC** | 1.86 ± 0.08 | 0.8 ± 0.1 |
| **StrM** | 1.26 ± 0.04 | 0.28 ± 0.09 |
| **CBU1838** | 0.87 ± 0.02 | 0.12 ± 0.03 |
| **AoEvaD** | 0.31 ± 0.01 | 0.28 ± 0.01 |
| **ChmJ** | 0.48 ± 0.07 | 0.2 ± 0.1 |

### Sugar backbone rearrangement is not observed with either StrM or CBU1838

In neither NMR nor GC-MS data was there any evidence of a rearrangement of the sugar backbone. The proposed biosynthetic pathways of both StrM and CBU1838 require such a rearrangement, catalysed by either the epimerase or a synthase/reductase for each pathway. Our direct detection of product formation and chemical structure analysis might have allowed us to detect this re-arrangement. However, as the five-membered rings are likely to be substantially disfavoured at equilibrium, the possibility remains that re-arranged structures were present but below the limit of detection.

### The structures of StrM and CBU1838 are consistent with their role as double epimerases

We determined the structures of StrM and CBU1838 by X-ray crystallography, in the presence and absence of TDP (Table S1). Both proteins show an overall dimeric architecture consistent with previously solved members of this family (Figure 6A). Both proteins show a strand exchange in common with orthologues. TDP binds to both proteins in a similar manner (Figure 6B,C). TDP is held in place by a network of amino acid side chains^a^ (R25^*^, F28^*^, E30^*^, Q48, R61, H64, Y140, K170) from both protomers (starred residues from the domain swapped strands).

**Figure 6:**
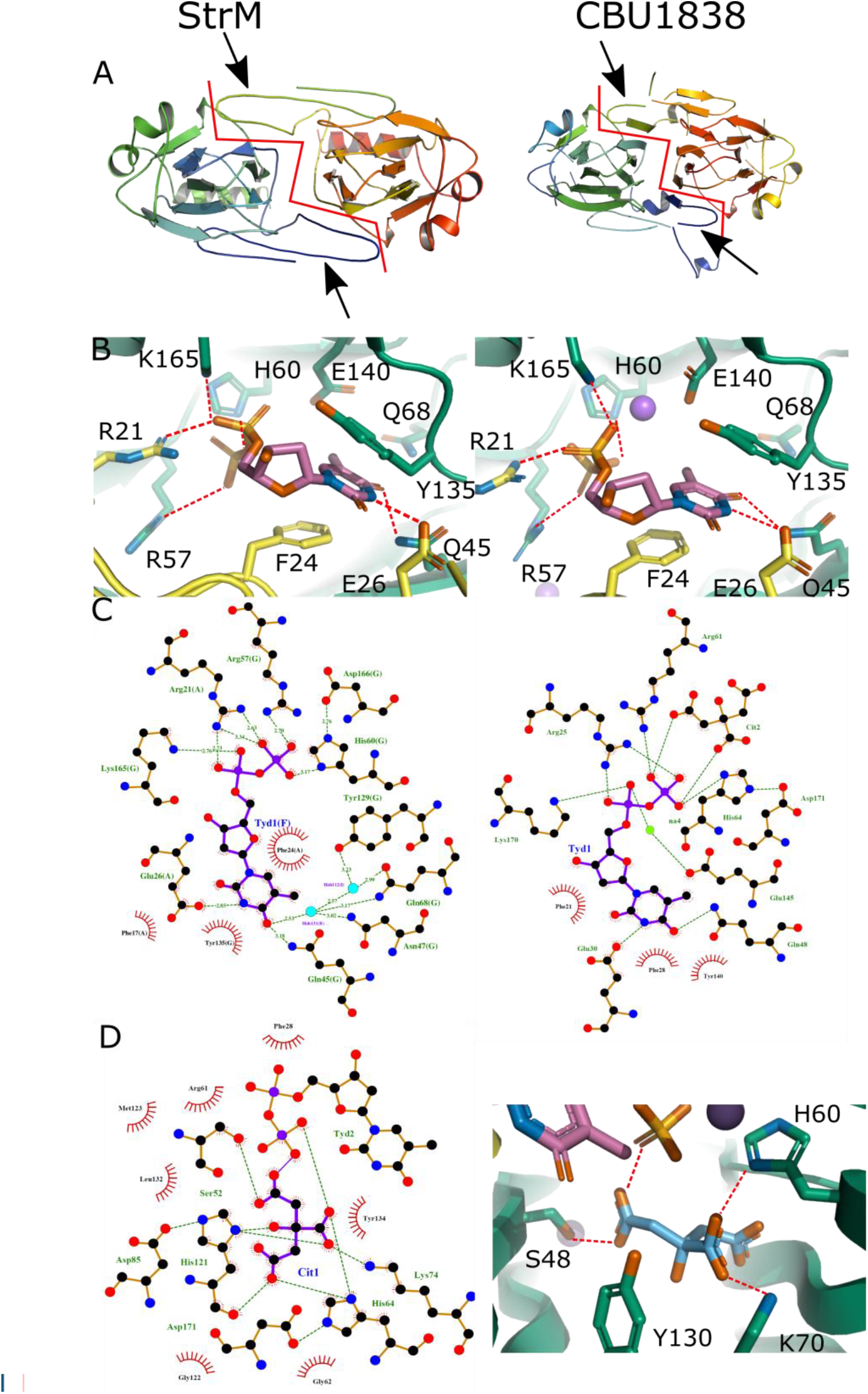
StrM and CBU1838 show expected interactions with ligands. **A:** Ribbon diagram of the structure of StrM (left) and CBU1838 (right). Both proteins form the domain-swapped dimer characteristic of the RmlC proteins. The red lines indicate the dimer interfaces. The domain-swapped strands are indicated by black arrowheads. Proteins are shown in cartoon format, coloured blue: protomer 1, N-terminus to red: protomer 2, C-terminus. **B,C:** Structures of both proteins complexed to TDP show a network of interactions conserved with other RmlC enzymes. Interaction maps for TDP with both enzymes were calculated using LigPlus v. 1.4.5. Structures are shown with protein backbone shown as cartoon, with TDP and interacting residues shown as sticks. Colours: nitrogen, blue; oxygen, red; phosphorus, orange; protomer 1 carbon, green; protomer 2 carbon, yellow; TDP carbon, rose. Hydrogen bonds are shown as red dashes. **D**: A citrate molecule occupies the substrate sugar site and makes interactions with key residues. Left: LigPlus summary of interactions. Right: Structural representation of the citrate. The two conserved catalytic residues H60 and Y130 flank the citrate. Representation as in B, with citrate carbons shown in skyblue. Panels A, B and D made using PyMOL v. 2.3.4.

Additional interactions are formed through bound solvent from N50, Q72, Y134 and E145. These are all strongly conserved across RmlC orthologues (Figure 2). The CBU1838-TDP structure showed a citrate molecule (carried over from crystallisation) bound to the active site. This molecule occupies the same space as the substrate in previously solved structures. The citrate interacts with S52, H64, K74 and Y134 from the protein, and TDP (Figure 6D). These amino acids are well conserved amongst orthologues, and H64/Y134 are the catalytic base and acid proposed from previous studies. The conformation of these side chains is very similar to that seen in complexes of other enzymes with substrate analogues (Figure S6). These structures give further confidence that CBU1838 and StrM are indeed orthologues of RmlC. They have a very similar fold, show clear and specific binding to TDP, and retain the key catalytic residues in the correct orientation.

## Discussion

Streptose and DHHS are monosaccharides produced by a single organism each. Both sugars are proposed to be synthesised through modified TDP-rhamnose pathways. *S. griseus* and *C. burnetii* contain putative orthologues of this biosynthetic pathway. Both organisms contain a single TDP-6-deoxy-D-*xylo*-4-hexulose 3’’,5’’-epimerase orthologue (StrM and CBU1838). We sought to determine whether these enzymes perform both epimerisations, and whether either might also perform the sugar ring rearrangement activity. Answering these questions will provide further insight into the biosynthesis of these unusual sugars.

CBU1838 showed robust double epimerase activity both in vitro and in vivo. CBU1838 complemented an *rfbC* (*rmlC*) mutation, restoring flux through the pathway and consuming cellular TDP-glucose and TDP-6-deoxy-D-*xylo*-4-hexulose (Figure 3). CBU1838 and StrM were competent to replace RmlC in vitro in a biosynthetic pathway (Figure 4). It was notable that substantially more CBU1838 was required to restore activity than RmlC or StrM. Similarly, CBU1838 showed a lower *k*_*cat*_, and higher *K*_*M*_, than either RmlC or StrM (Table 2). This is consistent with the proposed role of CBU1838, as the DHHS pathway requires a substrate with a 6’’-hydroxyl. CBU1838 contains a methionine residue (M123) in place of a phenylalanine conserved among other RmlC orthologues, which would provide space for the hydroxyl. The *k*_*cat*_ and *K*_*M*_ values that we obtained for StrM and RmlC are in the same order as those observed for other RmlC paralogues (Table S2). It is notable that our data were collected at 37 °C, whilst the comparative data were collected at 21-25 °C. It is likely that other enzymes would react faster at higher temperatures but might show a higher *K*_*M*_^55^.

Our NMR data show that CBU1838 and StrM have similar activity to *E. coli* RmlC, with CBU1838 again showing a reduced comparative activity. The rate of proton exchange at the 3’’ position was faster than that at the 5’’ position for all three enzymes (Figure 5; Table 3). The *M. tuberculosis* RmlC showed faster epimerisation at the 5’’ position ^41^. Dong et al. suggested that in the likely catalytic conformation, the 5’’ proton is more acidic than the 3’’ proton ^41^. Our experiments measured the loss of proton signal by NMR, whilst the previous study measured the product masses by GC-MS. Furthermore, our study used chemically prepared TDP-6-deoxy-D-*xylo*-4-hexulose, which has only recently become commercially available. This may explain some differences between our study and previous work. We note that as all the reactions involve proton transfers at all steps ^40^, there may be a strong kinetic isotope effect ^56^ on either or both epimerisations. As our methods for differentially measuring the two epimerisations rely on deuterium, this confounding effect cannot be excluded.

The structures of CBU1838 and StrM are highly similar to previously solved structures of paralogues from other species. TDP is held in place by residues that are strongly conserved (Figure 2, 6) ^43,46,49^. Although we were not able to determine a structure with a TDP-linked sugar, the side chains lining the sugar binding (and catalytic) cavity adopt conformations that are highly similar to those in previous structures ^38,41,43-48,50^. The proposed catalytic residues, H60 and Y130, occupy a very similar space to the equivalent residues in substrate analogue co-crystal structures ^41,50^. As their local hydrogen-bonding networks are also conserved, it is likely that they will also be catalytic in these proteins. The presence and conformation of these catalytic residues cannot predict the specificity of the enzymes. Whilst one example of a 5’’-specific enzyme showed a mutation that altered the conformation of an important active site side chain ^47^, 3’’-specific epimerases have shown a complete complement of active site residues ^44,48^. Our development of a functional assay in *E. coli* allowed us to confirm the activity of StrM and CBU1838 *in vivo* (Figure 3); although the 5’’-specific enzyme EvaD was expressed to the same levels as the active enzymes, it was not competent to complement *rmlC*. However, this assay did not distinguish between StrM (likely a direct RmlC orthologue) and CBU1838 (which is expected to use the 6’’-hydroxylated analogue).

The evidence presented in this and previous studies strongly suggests that both CBU1838 and StrM act as TDP-4-keto-(6-deoxy)-glucose 3’’, 5’’ double epimerases. The lower rates observed for CBU1838 are consistent with its natural substrate retaining the 6’’-hydroxyl group; an activity to generate this substrate has not yet been identified. This suggests that the streptose and DHHS synthase enzymes (Figure 1) accept a substrate that is epimerised at both the 3’’ and 5’’ positions. Although our data suggest that these enzymes cause proton exchange at the 3’’ carbon more efficiently than the 5’’ carbon, we cannot exclude that the kinetic isotope effect could be confounding. Our results provide powerful evidence that DHHS biosynthesis in *Coxiella* has convergently evolved to use a similar pathway to streptose biosynthesis in *S. griseus*. The key challenge that this presents is to identify the enzymes that provide the two remaining activities to generate DHHS.

## Methods

### Gene cloning and synthesis

The gene fragments for *rmlB, rmlC* and *rmlD* were amplified from *E. coli* DH5α (NEB) genomic DNA (gDNA) by PCR. The *CBU1838* gene fragment was codon-optimised using IDT’s tools and ordered as a gBlock from IDT and amplified by PCR. Primer sequences are provided in Table S3. Gene fragments were cloned into pNIC28-Bsa4 (Addgene #26103, gift of Opher Gileadi, SGC Oxford) using ligation-independent cloning (LIC), following published methods ^57^. Briefly, 16.5 µg pNIC28-Bsa4 plasmid was digested with 60 U *Bsa*I (NEB) in 100 µL following the manufacturer’s protocol, and incubated for 2 h at 50 °C. To generate the LIC cohesive ends in both the plasmid and PCR product, a mixture of 1 µL water, 5 µL *Bsa*I-digested plasmid/PCR-pure DNA, 2 µL 5X T4 DNA polymerase buffer (Fermentas), 1 µL 25 mM dGTP (insert) or dCTP (plasmid), 0.5 µL 100 mM DTT, 0.5 µL T4 DNA polymerase (Fermentas) was prepared, and incubated at 22 °C for 30 min, then 20 min at 75 °C. 2 µL of treated DNA product was added to 1 µL of treated plasmid. The mixture was incubated at room temperature for 10 min followed by transformation into 5-alpha competent cells (NEB). The *E. coli* RmlD contains an inactivating point mutation. This was corrected by site-directed mutagenesis (SDM) using the QuikChange Lightening kit (Agilent), following the manufacturer’s instructions.

TDP-sugar epimerase genes for *Streptomyces griseus strM, Amycolatopsis orientalis evaD* and *Streptomyces bikiniensis chmJ* were codon optimised using an in-house Python script (https://github.com/njharmer/CodonOptimise) and cloned into pET28b with an additional SUMO tag, pET28b, and pET21a respectively by Twist Bioscience. Plasmid were transformed into the expression strain *E. coli* BL21 (DE3) (Novagen) using kanamycin (50 µg/mL) for selection. For complementation of *E. coli rfbC* mutants, TDP-sugar epimerase genes for *strM, evaD* and *CBU1838* were cloned into the cloning vector pTWIST-A-MC under the control of the lac operon, with a strong ribosome binding site calculated by the RBS Calculator (De Novo DNA). The Genbank files for all plasmids are available in Supplementary Data.

### Preparation of electrocompetent *E. coli* mutant cell lines and complementation

*ΔrfbC E. coli* strains (Keio collection; Horizon Discovery #OEC4987-200827372) were grown overnight on LB plates with 50 µg/mL kanamycin. Colonies were inoculated from LB plates into 10 mL fresh LB broth and incubated overnight at 37°C with shaking at 225 rpm. 1 mL of overnight culture was inoculated into 100 mL fresh LB medium with kanamycin and incubated at 37 °C with shaking at 225 rpm until OD600 was approximately 0.6. Cells were harvested by centrifugation (4200 x*g*, 10 min, 4 °C). The cells were resuspended in 40 mL pre-chilled autoclaved Milli-Q water. Cells were harvested by centrifugation as above and resuspended in 40 mL pre-chilled autoclaved 10 % (v/v) glycerol. A minimum of three 10 % glycerol washes were conducted before cells were resuspended in 500 µL 10 % (v/v) glycerol.

Complementation plasmids (40 ng) were added to 50 µL electrocompetent cells, vortexed for 30 s and incubated on ice for 10 min. The suspension was transferred to a pre-chilled 0.1 cm electroporation cuvette (Thermo Scientific #5510-11). A single pulse of 1.8 kV was given by a Gene Pulser Xcell^®^ Microbial System (Bio-Rad #1652662). Cells were recovered by adding 1000 µL pre-warmed LB medium. The complemented cell suspension was incubated at 37 °C for 1 hour shaking at 225 rpm. Transformed cells were grown on LB agar plates supplemented with 100μg/mL ampicillin overnight at 37 °C.

### Detecting expression of complemented *E. coli*

Complemented *E. coli* strain colonies were inoculated into 100 mL bottles containing 10 mL LB medium, 100 μg/mL ampicillin and incubated overnight at 37 °C with shaking at 225 rpm. 1 mL of each overnight culture was inoculated into individual 500 mL conical flasks containing 100 mL LB medium supplemented with ampicillin and incubated at 37 °C with shaking at 225 rpm. When the OD_600 nm_ reached 0.6, gene expression was induced by adding 200 µM IPTG. Cultures were incubated at 20 °C with shaking at 225 rpm overnight. 50 mL aliquots of overnight cultures were harvested by centrifugation (4500 x*g*, 30 min, 4 °C). Cells were lysed using 1 mL BugBuster Master Mix (Millipore #71456-4), following the manufacturer’s instructions. The soluble and insoluble fractions were separated by centrifugation at 16 000 x*g* for 30 min at 4 °C. Samples were separated on a 4-12% ExpressPlus™ PAGE gel (GenScript #M41212) following the manufacturer’s instructions. Samples were transferred to a nitrocellulose membrane (Sartorius Stedim Biotech #11327-41BL) using a Pierce G2 Fast Blotter (Thermo Scientific # 15146375) using the pre-programmed protocol. Antibodies were added using an iBind Western device (ThermoFisher #SLF1000), following the manufacturer’s recommendations. The primary and secondary antibodies used were anti-penta His (Qiagen #34660) and goat anti-mouse IgG (LiCOR #926-68070) at a dilution of 1:1000. Blots were imaged using an Odyssey Clx imaging system (LiCOR).

### Sample preparation for LC-MS QQQ analysis

Complemented *E. coli* strain colonies were inoculated into 100 mL bottles containing 10 mL LB medium, 100 μg/mL ampicillin for complemented strains, and incubated overnight at 37 °C with shaking at 225 rpm. 1 mL of each overnight culture was inoculated into individual 500 mL conical flasks containing 100 mL LB medium supplemented with ampicillin as necessary and incubated at 37 °C with shaking at 225 rpm until OD_600 nm_ reached 0.6. Expression was induced by adding 200 µM IPTG. Cultures were incubated at 20 °C with shaking at 225 rpm overnight. 50 mL aliquots of overnight cultures were harvested by centrifugation (4500 x*g*, 30 min, 4 °C). Cellular contents were extracted into 1 mL 50 % (v/v) acetonitrile. Cell suspensions were incubated at 25 °C for 30 min and centrifuged at 20 000 x*g* for 30 min at 4 °C. The supernatant was filtered using Millex Non-Sterile Low Protein Binding Hydrophilic LCR (PTFE) Membrane (0.45 µm) filters (Merck #SLLHR04NL) into 1.5 mL MS vials with silicone/PTFE septa and stored at -20 °C for later analysis.

### Quantitative LC-MS QQQ Analysis

Quantitative analysis was performed using an Agilent 6420B triple quadrupole (QQQ) mass spectrometer (Agilent Technologies, Palo Alto, USA) coupled to a 1200 series Rapid Resolution HPLC system. 5 µL of sample extract was loaded onto an Agilent Poroshell 120 HILIC-Z, 2.7 µm, 2.1 × 150 mm analytical column (Agilent Technologies #673775-924). For detection using negative ion mode, mobile phase A comprised 90 % (v/v) LC-MS grade acetonitrile with 10 mM ammonium acetate and 5 μM medronic acid, and mobile phase B was 100% water (LC-MS grade) also with 10 mM ammonium acetate and 5 μM medronic acid. The following gradient was used: 0 min – 10% B; 6 min – 35% B; 10 – 13 min – 40% B; 15 min – 10% B; followed by 5 minutes re-equilibration time at a flow rate of 0.25 mL min^-1^ with the column held at 25 °C for the duration. The QQQ source conditions for electrospray ionisation were: 350 °C gas temperature with a drying gas flow rate of 11 L min^-1^ and a nebuliser pressure of 35 psig. The capillary voltage was 4 kV.

### MS Data Analysis

Data analysis was undertaken using Agilent MassHunter Quantitative Analysis software (version B.07.01, SP1). Data was normalised to the internal standard, ^13^C TDP-glucose, and the optical density of each culture pre-extraction. TDP-glucose, TDP-rhamnose, GDP-mannose, GDP-fucose and UDP-glucose concentrations were calculated using standard calibration curves. For the other compounds (where a standard was not available) normalised peak areas were compared.

### Expression and purification of TDP-sugar epimerases

RmlB, RmlC, RmlD, StrM, EvaD and ChmJ were expressed in 500 mL of ZYM-5052 auto-induction media supplemented with 100 µg/mL kanamycin following the methods of ^58^. Each flask was inoculated with 10 mL of an overnight culture and grown at 37 °C with agitation at 200 rpm until OD_600_ reached 0.6. Cultures were further incubated at 20 °C for 18 hours with agitation. Cells were harvested by centrifugation at 4500x*g* for 30 min at 4 °C. The pellet was resuspended in binding buffer (20 mM Tris-HCl, 500 mM NaCl, 10 mM imidazole, pH 8.0) and lysed by sonication (SONIC Vibra cell VCX130). The sample was clarified by centrifugation (24 000 x*g* for 30 min at 4 °C). The soluble fraction was purified using an ÄKTAxpress chromatography system (Cytiva). The sample was purified using a 1 mL HisTrap crude column (Cytiva). After loading sample, the column was washed with binding buffer, and the protein step eluted into binding buffer supplemented with 250 mM. The eluate was purified over a Superdex 200 pg 16/600 size-exclusion column (Cytiva) and eluted isocratically into 10 mM HEPES, 500 mM NaCl, pH 7.5 (for coupled assays and crystallisation) or 10 mM Na_x_HPO_4_, 500 mM NaCl, pH 7.5 for NMR studies. The eluted protein was concentrated using a Vivaspin centrifugal concentrator (Generon) and stored at -20 °C with 20 % (v/v) glycerol for enzymatic assays; or stored at -80°C in small aliquots without any glycerol for crystallization or NMR.

### Biochemical assays

The TDP-rhamnose biosynthesis assay coupled the product of RmlC paralogues to RmlD. In brief, reactions were performed in 96-well flat-bottomed plates (Greiner #655001) in a total volume of 200 µL. For initial studies, reactions consisted of 50 mM HEPES pH 7.5, 20 mM MgCl_2_, 350 µM NADPH, 4 µM NAD^+^, 0.4 µM RmlB, 2 µM RmlC paralogue and 2 µM RmlD. Reactions were initiated by addition of 500 µM TDP-glucose (Carbosynth #MT04383). Reactions were monitored by measurement of the absorbance at 340 nm over 30 min in an Infinite M200PRO plate reader (Tecan) incubating at 30 °C. For determination of kinetic parameters, RmlB and NAD^+^ were removed and TDP-glucose was replaced with TDP-6-deoxy-D-*xylo*-4-hexulose (Carbosynth #NT29846) added at a concentration range between 0-3 mM (*Ec*RmlC / *Sg*StrM) or 0-9 mM (CBU1838 or *Ao*EvaD). Three experimental replicates were performed for all reactions. Data were fitted to the Michaelis-Menten equation using Graphpad v. 8.1.2.

### Deuterium-incorporation analysis by ^1^H NMR

Reaction mixtures contained 10 mM MgCl_2_, 2.38 mM TDP-6-deoxy-D-*xylo*-4-hexulose (Carbosynth) and 8 µM NAD^+^ in a final volume of 250 µL deuterated 100 mM phosphate buffer pD 7.0. A “zero time” spectrum was recorded without any epimerase. Epimerases (38 nM *Ec*RmlC, 46 nM *Sg*StrM, 177 nM CBU1838, or 530 nM *Ao*EvaD) were added to the above reaction mixture to a final volume of 250 µL.

All data were collected on a Bruker Neo 600 MHz spectrometer equipped with TCI cryoprobe. Standard Bruker proton experiments were carried out at 298 K (25 °C). Periodic ^1^H NMR experiments (64 scans) were carried out in 3 mm NMR tubes. ^1^H NMR spectra were recorded every 10 min over a 90 min period, set using the Topspin multi_zgvd command. Later intervals were set manually to record at 2 h, 4 h, 8 h, 18 h and 24 h, as appropriate. TopSpin 4.0.6 was used for data processing using command ‘efp’ followed by automatic phase correction (‘apk’) and automatic baseline correction (‘abs’). The residual water peak at δ4.7058 ppm was used as reference for spectra calibration. A set of selected diagnostic signals from no-enzyme control samples was integrated for calibration of the relative abundance of each signal. An integral of H-6 thymidine peak (^1^H) was used for peak intensity calibration of the rest of integrals of diagnostic signals and GraphPad Prism v8.1.2 was used to analyse the data obtained.

### Deuterium-incorporation analysis GC-MS

0.5 mg TDP-6-deoxy-D-*xylo*-4-hexulose (Carbosynth) was incubated in deuterated buffer alone, or with the addition of either 1 µM of EcRmlB, EcRmlC, SgStrM, CBU_1838 or AoEvaD in 250 µL final volume. After 18 h, enzymes were inactivated and removal by ethanol precipitation followed by centrifugation at 14,000 xg for 10 min. Sodium borohydride (∼2 mg) was added to each sample for the reduction of keto-moeity followed by incubation at room temperature for 2 h, with shaking at 120 rpm. The reaction was terminated with few drops of glacial acetic acid, followed by 20 µL 10 % (v/v) acetic acid in methanol. Solvents were evaporated under reduced pressure using GeneVac, for 1 h at 35 °C and 25 µg myo-inositol was added to the sample as an internal standard. The sugar-nucleotide bond was hydrolysed by using 250 µL 2 M trifluoroacetic acid (TFA) followed by incubation at 100 °C for 1.5 h. After cooling the samples, solvents were evaporated again and 100 µL isopropanol was added before a final evaporation in order to remove traces of TFA for 1 h.

A second reduction was achieved by addition of 250 µL 1 M ammonia, containing 10 mg/mL sodium borohydride for 1 h and then 250 µL acetic acid-methanol (1:9, v/v) was added. The samples were dried under reduced pressure (GeneVac, 1h at 40 °C) and the addition of acetic acid-methanol followed by drying was repeated once again. This was followed by addition of 250 µL methanol (without acetic acid) twice with evaporating the solvent off each time (20 min each). The resulting sample was acetylated by incubation at 120 °C for 20 min in a mixture of 100 µL acetic anhydride and 100 µL pyridine. After this, approximately 200 µL toluene was added, and the mixture was concentrated under reduced pressure. The addition of toluene and evaporation was repeated at room temperature.

The alditol acetates were separated by partition between equal volume of methylene chloride and water. After partition the methylene chloride layer was left to evaporate slowly at room temperature. Finally, samples were dissolved in acetone and analysed by gas chromatography-mass spectrometry (GC-MS).

### Crystallization

Protein concentrations used for crystallization were 5.85 mg/mL (*Sg*StrM) and 5.2 mg/mL or 7.8 mg/mL (CBU_1838). Crystals were grown using the microbatch method using an Oryx8 crystallization robot (Douglas Instruments). Crystals were grown in hydrophobic plates (Douglas Instruments #VB-Silver-1/1) and covered in a 1:1 mix of paraffin oil and silicone oil (Molecular Dimensions). Crystals were grown either as a 1:1 mixture of protein and mother liquor, or as a 3:2:1 mixture of protein, mother liquor and seed stocks where seeds had been prepared from previous crystals. The successful co-crystallisation conditions, substrates soaking conditions, and cryoprotectants used are detailed in Table S4.

### X-ray data collection and structure determination

Data were collected at Diamond Light Source (Didcot, UK) at 100 K using Pilatus 6M-F detectors (Table S1). All data were processed using XDS ^59^. CBU1838 data were subjected to anisotropic ellipsoidal truncation using the STARANISO server ^60^. The resolution limit for StrM datasets was set at a point where CC_1/2_ was above 0.3 in the highest resolution shell. CBU1838 data were processed to include regions where the mean I/σ was systematically above 1.2. Further data processing and structural studies was carried out using the CCP4 program package ^61^ and the CCP4i2 interface ^62^. Molecular replacement was used to provide initial phases for datasets using Morda ^63^ and MolRep ^64^. Density fitting was performed using Coot ^65^ and refined with REFMAC5 ^66^. Model validation was carried out with internal modules of CCP4i2 and Coot, employing MolProbity calculations ^67^.

## Acknowledgements

The authors thank Simone de Rose (University of Exeter) for assistance with crystallisation.

SR, MVV, RF and NJH were funded by BBSRC grant BB/N001591/1. ARC and NJH were funded by BBSRC grant BB/M016404/1 and by Dstl grant DSTLX-1000098217. Work at the John Innes Centre (MR, SAN, RAF) was supported by the UK BBSRC Institute Strategic Program on Molecules from Nature -Products and Pathways [BBS/E/J/000PR9790] and the John Innes Foundation, and the InnovateUK: IBCatalyst (GrantBB/M02903411).

## Author contributions

RAF, JP and NJH designed the study and obtained funding. ARC, SR, MVV, MR, SAN, MC and DS collected data. ARC, SR, MVV, MR, MC, DS, MI, RAF and NJH analysed data. ARC and NJH wrote the initial manuscript. All authors contributed to revision of the manuscript.

## Footnotes

a Residue numbers given are for CBU1838. The equivalent residues in StrM are (in order mentioned on this page) R21^*^, F24^*^, E26^*^, Q45, R57, H60, Y135, and K165; N47, Q68, Y129, and E140; S49 and K70.

