## Supplemental Figures and Tables for "Spinning sugars in antigen biosynthesis: a direct study of the *Coxiella burnetii* and *Streptomyces griseus* TDP-sugar epimerases"

**Supplementary information**

**Table 1: Crystallographic data table**

| Protein complex | CBU1838 xylose | CBU1838 TDP | StrM apo | StrM TDP |
|---|---|---|---|---|
| Data collection statistics | | | | |
| Beamline | I04 Diamond | I04 Diamond | I03 Diamond | I04-1 Diamond |
| Wavelength (Å) | 0.9795 | 0.9795 | 0.9763 | 0.9159 |
| Space group | C2 | $C222_1$ | $C222_1$ | $C222_1$ |
| Unit Cell Parameters a, b, c (Å) | 77.0, 84.6, 77.7 | 78.9, 82.2, 163.8 | 42.3, 132.4, 77.6 | 42.4, 131.5, 78.5 |
| $\alpha$, $\beta$, $\gamma$ (°) | 90.0, 113.6, 90.0 | 90.0, 90.0, 90.0 | 90.0, 90.0, 90.0 | 90.0, 90.0, 90.0 |
| Resolution range (Å)$^{a}$ | 51.39 – 1.44 (1.61-1.45) | 81.92-1.87 (2.0-1.87) | 66.18 – 1.34 (1.43 - 1.34) | 50.42-1.90 (1.94-1.90) |
| Total reflections $^{a}$ | 182,903 (7,987) | 247,730 (12,776) | 230,889 (11,075) | 76,059 (4,895) |
| Unique reflections $^{a}$ | 54,839 (2,742) | 37,452 (1,874) | 36,479 (1,824) | 17,624 (1,122) |
| Completeness spherical (%)$^{a}$ | 65.9 (11.6)$^{b}$ | 84.8 (25.0)$^{b}$ | 74.0 (20.8)$^{b}$ | 99.4 (99.6) |
| Multiplicity $^{a}$ | 3.3 (2.9) | 6.6 (6.8) | 6.3 (6.1) | 4.3 (4.4) |
| $R_{meas}$ (%) $^{a,d}$ | 5.7 (85.2) | 8.5 (140.0) | 14.2 (249.5) | 24.5 (240.5) |
| $\langle I\rangle/\langle\sigma(I)\rangle$ $^{a}$ | 12.5 (1.5) | 12.5 (1.3) | 5.6 (0.3) | 5.5 (0.7) |
| $CC_{1/2}$ $^{a,c}$ | 0.999 (0.585) | 0.998 (0.67) | 0.996 (0.304) | 0.991 (0.312) |
| Wilson B-factor $^{e}$ (Å$^{2}$) | 27.9 | 50.7 | 17.9 | 30.7 |
| Refinement statistics | | | | |
| $R_{work}$ | 0.168 | 0.209 | 0.201 | 0.203 |
| $R_{free}$ | 0.199 | 0.237 | 0.234 | 0.249 |
| No. of protomers in a.u. | 2 | 2 | 1 | 1 |
| Number of atoms | | | | |
| Protein | 3,540 | 3,324 | 1,727 | 1648 |
| Ligands | 53 | 100 | - | 25 |
| Solvent | 459 | 292 | 228 | 180 |
| Number of protein residues | 390 | 380 | 200 | 203 |
| RMS bond lengths (Å) | 0.008 | 0.011 | 0.009 | 0.006 |
| RMS bond angles (°) | 1.45 | 1.59 | 1.81 | 1.40 |

| | | | | |
|---|---|---|---|---|
| Ramachandran favoured (%)[f] | 99.48 | 97.61 | 97.98 | 98.51 |
| Ramachandran outliers (%)[f] | 0.0 | 0.27 | 0.0 | 0.0 |
| Clashscore [e] | 9.5 | 7.9 | 6.6 | 5.94 |
| Average B-factor protein (Å$^2$) | 22.3 | 41.2 | 13.9 | 26.8 |
| Average B-factor ligands (Å$^2$) | 30.1 | 75.5 | - | 52.4 |
| Average B-factor solvent (Å$^2$) | 35.2 | 48.5 | 26.6 | 36.7 |
| RCBS PDB code | 7PVI | 7PWB | 7PWI | 7PWH |

[a]Values for the highest resolution shell are given in parentheses.

[b] Values are given for data subjected to anisotropic ellipsoidal truncation using the STARANISO server [1]

[c] $R_{meas} = \Sigma_h[m/(m - 1)]^{1/2} \Sigma_i|I_{h,i} - <I_h>|/ \Sigma_h \Sigma_i I_{h,i.}$

[d] $CC_{1/2}$ is defined in [2].

[e] Wilson B-factor was estimated by SFCHECK [3].

[f] The Ramachandran statistics and clashscore statistics were calculated using MOLPROBITY [4].

**Table S2: Kinetic data for RmlC paralogues from diverse species.**

| Species | $k_{cat}$ (s$^{-1}$) | $K_M$ for TDP-6-deoxy-D-*xylo*-4-hexulose (μM) | Temperature (° C) | References |
|---|---|---|---|---|
| *S. suis* | 10.4 ± 0.3 | 29 ± 3 | 21 °C | [5,6] |
| *S. enterica* serovar Typhimurium | 19.2 ± 0.5 | 81 ± 8 | 21 °C | [5,6] |
| | 39.0 ± 6.6 | 710 ± 170 | 25 °C | [7,8] |
| *M. tuberculosis* | 18.0 ± 0.9 | 211 ± 43 | 25 °C | [9] |
| *A. thermoaerophilus* | 2.0 ± 0.7 | 62 ± 6 | 25 °C | [8] |
| *E. coli* | 5.5 ± 0.1 | 830 ± 40 | 37 °C | This study |
| *S. griseus* StrM | 5.4 ± 0.1 | 600 ± 40 | 37 °C | This study |
| *C. burnetii* CBU1838 | 1.75 ± 0.05 | 2,000 ± 200 | 37 °C | This study |
| *A. orientalis* EvaD | 0.39 ± 0.02 | 632 ± 98 | 37 °C | This study |

**Table S3: Primer sequences.** Primers used in this study were purchased from IDT.

| Primer name | Sequence (5’-3’) |
|---|---|
| RmlB_F | TACTTCCAATCCATGAAAATACTTGTTACTGGTGGCGCAGG |
| RmlB_R | TATCCACCTTTACTGTTACTGGCGGCCCTCATAGTTCTGTTCAATCC |
| RmlC_F | TACTTCCAATCCATGAATGTGATTAGAACTGAAATTGAAGATGTGC |
| RmlC_R | TATCCACCTTTACTGTCATGCAATTAATTTTAATCTGATAAGC |
| RmlD_F | TACTTCCAATCCATGAATATCCTCCTTTTTGGCAAAACAGG |
| RmlD_R | TATCCACCTTTACTGTTAAATTGCTGTAGTCGTAAATAATTCATTGAGC |
| RmlD_insC_F | GCAAAGCAGGCATTCCCCTTGCACTCAACAAGCTC |
| RmlD_insC_R | GAGCTTGTTGAGTGCAAGGGGAATGCCTGCTTTGC |
| CBU1838_F | TACTTCCAATCCATGCCGTTTGAATTTCAAAAAATGCTC |
| CBU1838_R | TATCCACCTTTACTGTTAAGAGCCATGATACTGCGC |

**Table S4: Crystallisation and soaking conditions.**

| | Structures | | | |
|---|---|---|---|---|
| Complex | CBU1838 xylose | CBU1838 TDP | StrM apo | StrM TDP |
| RCSB PDB code | 7PVI | 7PWB | 7PWI | 7PWH |
| Protein stock concentration (mg/mL) | 4.5 (co-crystallised with 30 % (w/v) xylose) | 4.5 (co-crystallised with 10 mM TDP) | 5.85 | 5.85 |
| Precipitant mixture | 25 % (w/v) PEG 3350, 100 mM Bis-Tris pH 5.5 | 8 % (w/v) PEG 4000, 0.1 M sodium acetate pH 4.6 | 20 % (w/v) PEG 6000, 100 mM $CaCl_2$, 50 mM HEPES pH 7.0 | 20 % (w/v) PEG 6000, 100 mM $CaCl_2$, 50 mM HEPES pH 7.0 |
| Ratio of protein: precipitant: seeds | 3:2:1 (1:1000 seed stock) | 3:2:1 (1:100 seed stock) | 1:1 | 1:1 |
| Cryoprotectant solution | 100 mM $(NH_4)_2SO_4$, 8 % (w/w) PEG 8000, 30 % (w/v) xylose | 7 % (w/v) PEG 4000, 30 % (v/v) PEG 400, 5 mM TDP, 29 mM citrate pH 4.5 | 10 % (w/v) PEG 6,000, 30 % (v/v) PEG 300, 100 mM $CaCl_2$, 50 | 10 % (w/v) PEG 6,000, 30 % (v/v) PEG 300, 100 mM $CaCl_2$, 50 |

| | | | | |
|---|---|---|---|---|
| | | | mM Tris pH 7.0 | mM Tris pH 7.0, 10 mM TDP |
| Soaking time (min) | 1 | 1 | 1 | 6 |

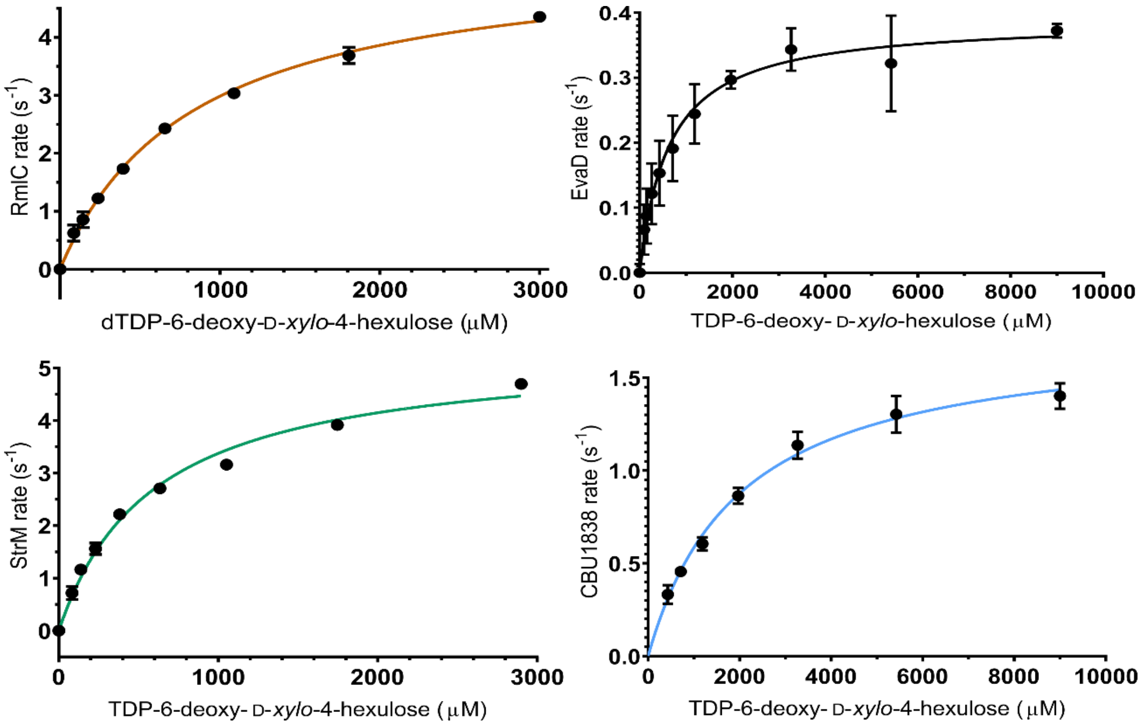


**Figure S1. Kinetic activity of TDP-sugar epimerases.**

RmlC and StrM were used at 0.2 µM, CBU1838 was used at 0.6 µM, and *Ao*EvaD at 1.0 µM, each with 0.8 µM RmlD, 350 µM NADPH, 20 mM $MgCl_2$, and 50 mM HEPES pH 7.0. Due to a slight lag phase, the steepest slope from each substrate concentration replicate was chosen for rate analysis. For RmlC and StrM at the highest substrate concentration, completion was not reached after 300-500 s. For CBU1838 completion was reached after 50-550 s, for *Ao*EvaD this was doubled. Data were fitted to the Michaelis-Menten non-linear regression analysis equation. Data were analysed using GraphPad Prism v8.1.2. Three experimental replicates were taken for each data point, and error bars represent the standard error in the mean.

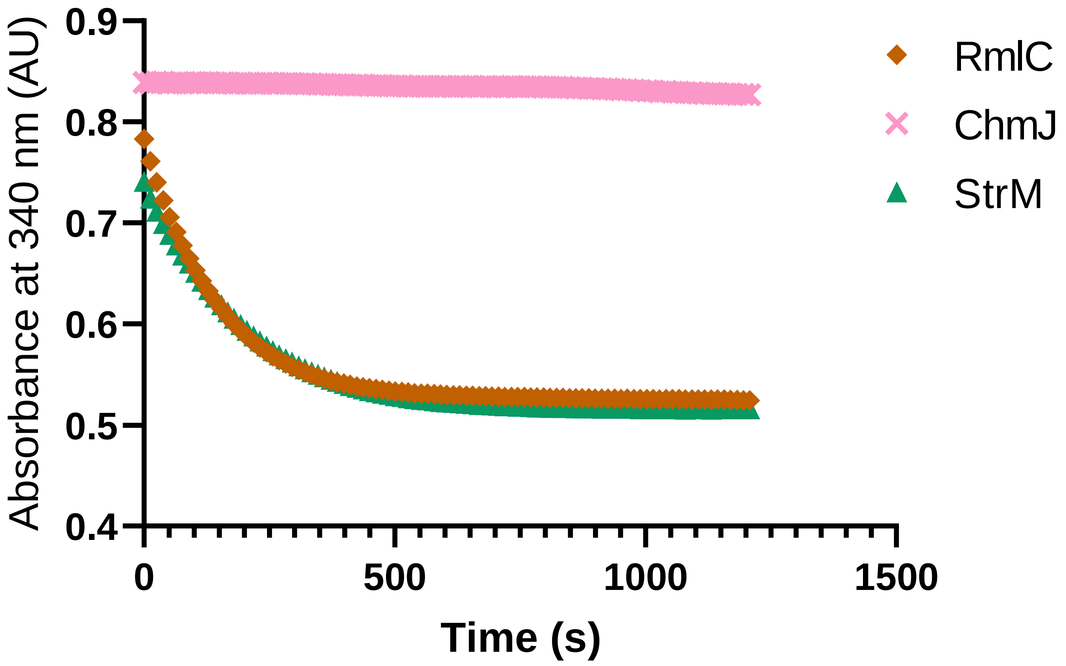


**Figure S2: ChmJ is not capable of 3'',5'' double epimerisation of TDP-6-deoxy-D-*xylo*-4-hexulose.** RmlC and StrM were used at 0.4 µM, whilst ChmJ was used at 1.6 µM, each with 0.4 µM each RmlB and RmlD, 5 µM $NAD^+$, 350 µM NADH, 500 µM $MgCl_2$, 150 µM TDP-glucose and 50 mM HEPES pH 7.5. Whilst RmlC and StrM consume all the TDP-sugar substrate, ChmJ does not cause any turnover. The observed rate is at the limit of detection of the assay ($10^{-5}$ AU $s^{-1}$), which is limited by the natural breakdown of NADPH at 37° C. Data were analysed using GraphPad Prism v8.1.2.

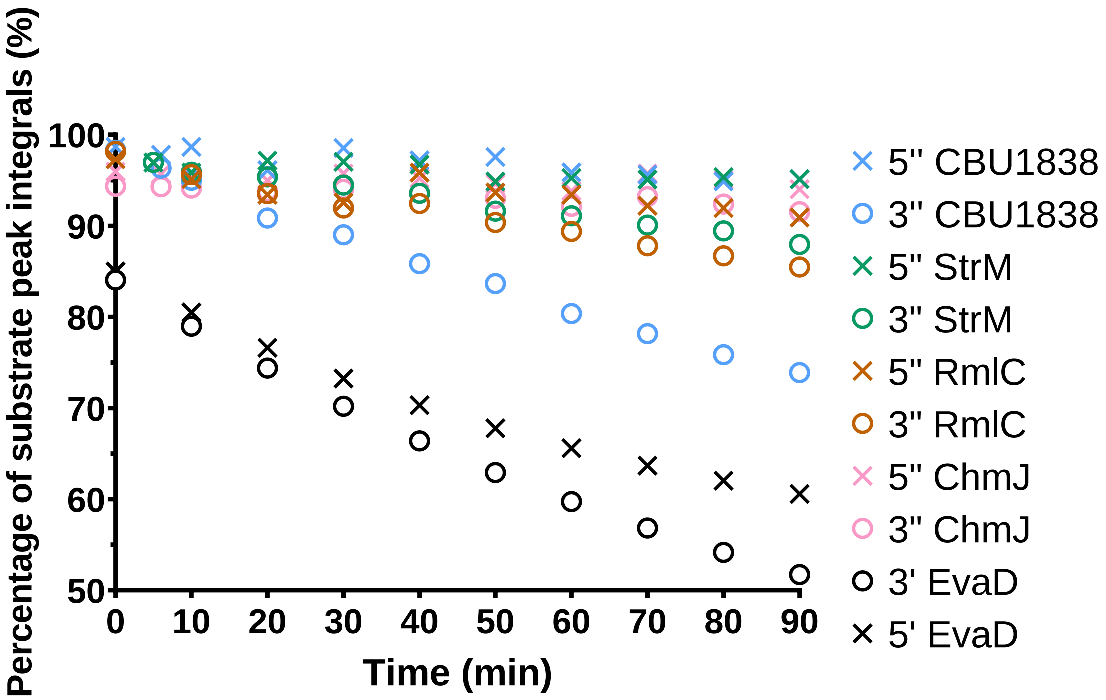


**Figure S3: Rate of RmlC paralogues monitored by NMR.** RmlC paralogues were incubated with purified TDP-6-deoxy-D-*xylo*-4-hexulose. 38 nM *Ec*RmlC, 46 nM *Sg*StrM, 177 nM CBU1838, or 530 nM *Ao*EvaD was used, with more enzyme used for the less active enzymes to ensure that a detectable rate was observed. The loss of proton signal at the 3'' and 5'' positions in the sugar was monitored by NMR. The rate was monitored every 5-10 minutes over 90 minutes to calculate an initial rate.

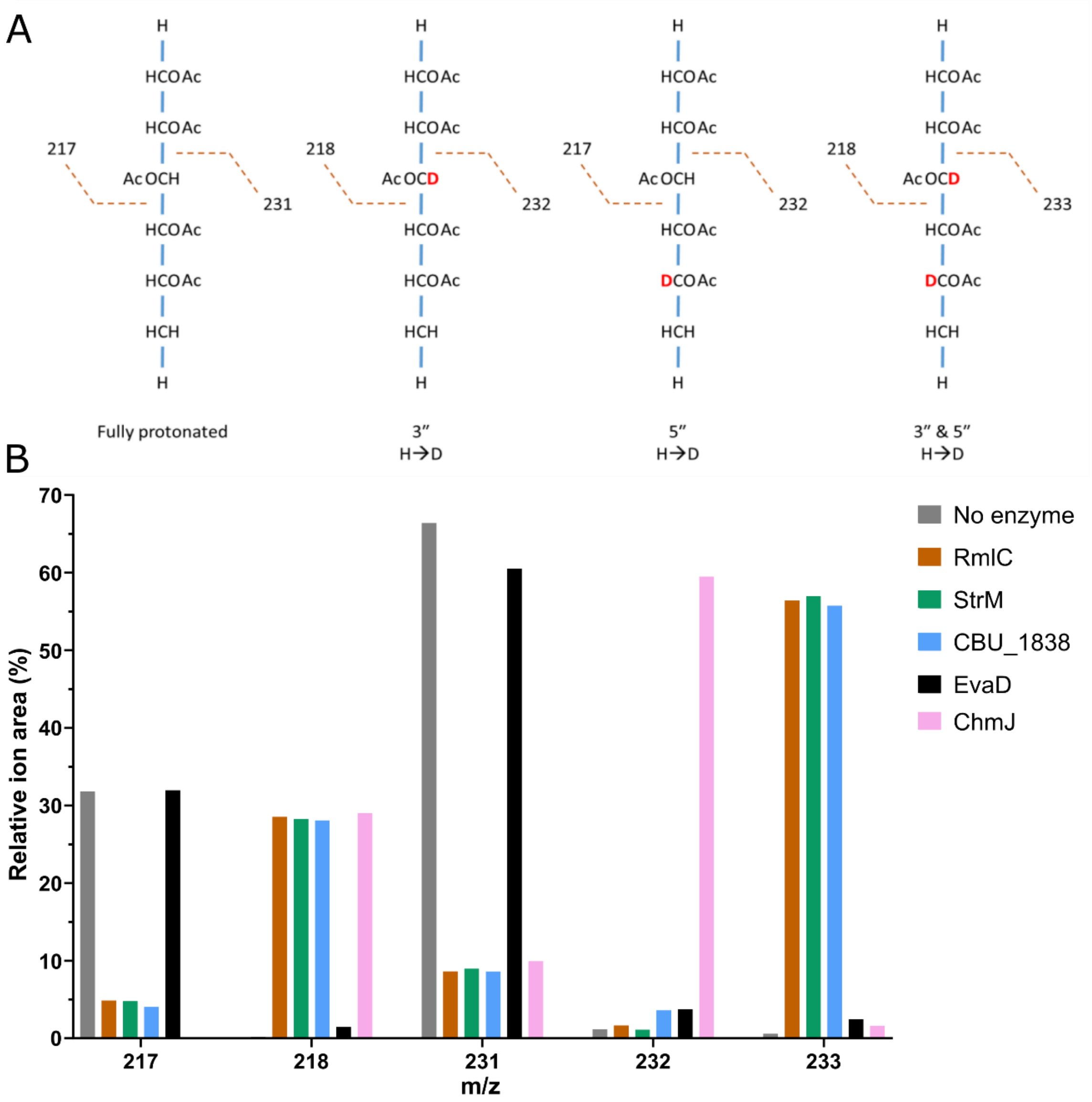


**Figure S4: Deuterium incorporation into TDP-6-deoxy-D-xylo-4-hexulose measured by GC-MS.** 0.5 mg TDP-6-deoxy-D-*xylo*-4-hexulose was incubated in deuterated buffer alone, or with the addition of 1 µM of *Ec*RmlC, *Sg*StrM, CBU1838, *Ao*EvaD or *Sb*ChmJ in 250 µL. After 18 h alditol acetates were prepared following the methods of [10-12]. **A.** The characteristic fragmentation products of the original sample and the mono/bi-substituted products. This highlights that the peak at m/z 233 is characteristic of the double substituted product; an ion pair at 218/232 m/z is characteristic of a 3'' single substitution; and an ion pair at 217/232 m/z is characteristic of a 5'' single substitution. **B**. Observed products after incubation with enzymes for 18 h. At the test concentrations, RmlC, StrM and CBU1838 show almost complete conversion to the 3'',5'' substituted product. EvaD shows a low level of substitution at both 3'' and 5'' positions. ChmJ shows almost complete substitution at the 3'' position, but little substitution at the 5'' position.

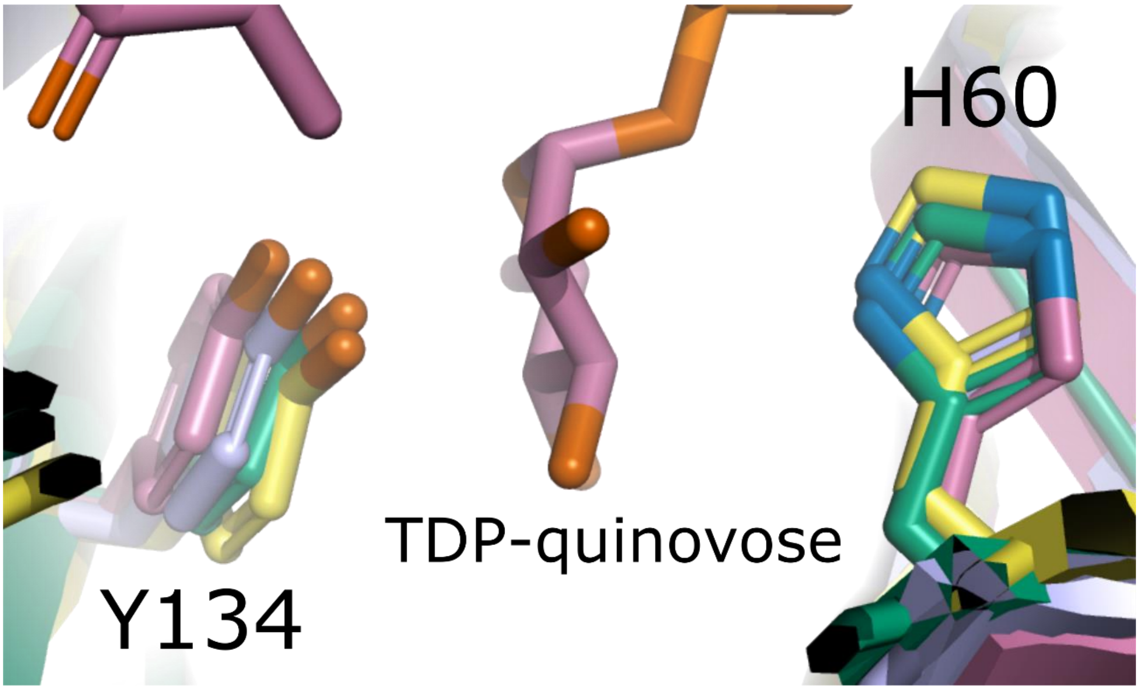


**Figure S6: Key StrM and CBU1838 catalytic residues retain the conformation observed in other paralogues.** The proposed catalytic acid and base Y134 and H60 are shown from the structures of CBU1838 (sea-green) and StrM (yellow) in comparison to the structures of *Mt*RmlC complexed to TDP-rhamnose (2IXC; light blue; [7]) and *Sb*ChmJ complexed to TDP-quinovose (4HMZ; rose; [13]). The TDP-quinovose and catalytic residues are shown as sticks; all other ligands have been removed. Protein backbones are shown as cartoons. Colours: nitrogen, blue; oxygen, red; phosphorus, orange. Structures were superimposed and the image generated using PyMOL v. 2.3.4.
